# FABP7 Progenitors are a Targetable Metabolic Root in the *BRCA1* Breast

**DOI:** 10.1101/2023.11.02.565360

**Authors:** Curtis W. McCloskey, Bowen Zhang, Matthew Waas, Golnaz Abazari, Foram Vyas, Kazeera Aliar, Pirashaanthy Tharmapalan, Abhijith Kuttanamkuzhi, Swami Narala, Jennifer Cruikshank, Stefan O.P. Hofer, Hartland W. Jackson, Thomas Kislinger, Hal K. Berman, Rama Khokha

**Affiliations:** Princess Margaret Cancer Centre, University Health Network, Toronto, Ontario, Canada; Department of Medical Biophysics, University of Toronto, Toronto, Ontario, Canada; Centre for Molecular and Systems Biology, Lunenfeld-Tanenbaum Research Institute, Mount Sinai Hospital, Toronto, Ontario, Canada; Department of Molecular Genetics, University of Toronto, Toronto, Ontario, Canada; Division of Plastic Surgery at the Department of Surgery and Department of Surgical Oncology, University Health Network, Toronto, Ontario, Canada; Department of Laboratory Medicine and Pathobiology, University of Toronto, Toronto, Ontario, Canada

## Abstract

It has been nearly 3 decades since the discovery of the *BRCA1/2* genes and their link to breast cancer risk, with prophylactic mastectomy remaining the primary management option for these high-risk mutation carriers. The current paucity of interception strategies is due to undefined, targetable cancer precursor populations in the high-risk breast. Despite known cellular alterations in the *BRCA1* breast, epithelial populations at the root of unwarranted cell state transitions remain unresolved. Here, we identify a root progenitor population that is dysregulated in *BRCA1* carriers stemming from the metabolic role of BRCA1. This fatty-acid binding protein 7 (FABP7) expressing luminal progenitor population is spatially confined to the mammary ducts, has enhanced clonogenic capacity, and is the predicted origin of mixed basal-luminal differentiation in the *BRCA1* but not *BRCA2* breast. We show global H3K27 acetylation is reduced within ductal FABP7 cells in *BRCA1* carriers *in situ*, linking to a non-canonical metabolic role of BRCA1 in regulating acetyl-CoA pools and *de novo* fatty acid synthesis. We demonstrate FABP7 progenitor capacity is preferentially ablated in *BRCA1* carriers through inhibition of fatty acid metabolism using an FDA-approved fatty acid synthase (FASN) inhibitor. This study lays the foundation for metabolic control of breast progenitor dynamics to mitigate breast cancer risk in the *BRCA1* breast.

---

Strides have been made in breast cancer risk stratification, yet major barriers remain in risk reduction for individuals facing up to a 75% lifetime risk and earlier onset of aggressive breast cancers. Recent efforts including the human breast cell atlas initiative are seeking to resolve the heterogeneity of the non-carrier and *BRCA1/2* mutation carrier human breast, across epithelial and stromal compartments^1–9^. These foundational studies have found stromal and immune drivers of high-risk phenotypes and highlight the need for targetable candidates in the epithelial compartment, which is known to harbor precursors for breast cancer subtypes.

We performed multiplexed single cell RNA-sequencing (MULTI-seq) on non-carrier and pathogenic *BRCA1/2* mutation carrier human breast tissues by mixing equal numbers of epithelial, endothelial, and fibroblast enriched cell fractions to encompass the cellular diversity of the human breast (**Figure 1A**). To mitigate biological flux over the menstrual cycle and the ovarian hormone-triggered expansion in mammary stem and progenitor pools^10–15^, we profiled follicular phase- staged samples. We applied bioinformatics tools designed to interrogate epithelial heterogeneity. Following quality control, 39049 cells from 11 patients (n=3 non-carrier, n=4 *BRCA1*, n=4 *BRCA2,* **Table S1**) were initially clustered and plotted using common UMAP coordinates revealing eleven known mammary cell types, including basal and luminal epithelial cells based on published lineage markers (**Figure S1A, S1B, Table S2**). Broadly, all cell types were represented in all patient samples (**Figure S1C**).

**Figure 1:**
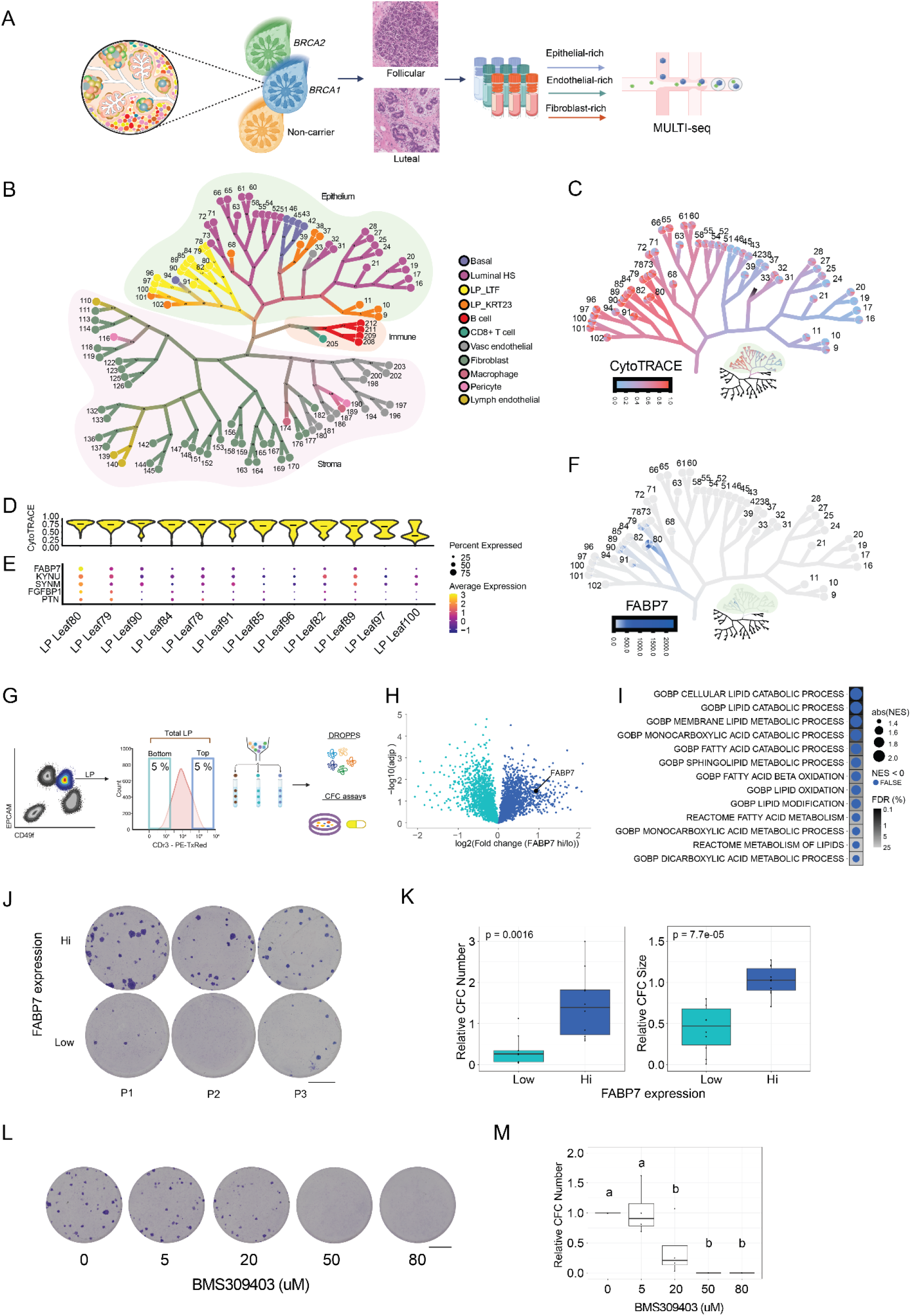
FABP7 marks a unique luminal progenitor subset. (A) Schematic of human breast workflow. Three cellular fractions from menstrual cycle staged non-carrier (n = 3), *BRCA1* (n = 4), and *BRCA2* (n = 4) human breast tissues were multiplexed for single-cell RNAseq using MULTI-seq. **(B)** TooManyCells (TMC) tree data abstraction annotated by broad cell type. PieCharts represent the proportion of each categorical variable within each leaf. Leaf (cluster) number is shown. **(C)** Epithelial subsetted TMC tree with CytoTRACE scores projected. **(D)** Violin plot of CytoTRACE scores per LP LTF mammary epithelial subset. Mean value per group is marked (black line). **(E)** Top 5 markers genes for CytoTRACE^hi^ LP leaf 80 are presented. **(F)** FABP7 expression projected onto epithelial subsetted TMC tree. **(G)** FACS workflow for the isolation of FABP7^hi^ (top 5 % CDr3), FABP7^lo^ (bottom 5 % CDr3), and total LPs for downstream Droplet-based one-pot preparation for proteomic samples (DROPPS) and colony-forming capacity (CFC) assays. **(H)** Volcano plot of differential proteome expression between FABP7^hi^ (dark blue) and FABP7^lo^ (teal) cells. **(I)** Pathway analysis of GO and Reactome terms enriched in FABP7^hi^ compared to FABP7^lo^ proteomes. **(J)** Representative CFC images from sorted FABP7^hi^ and FABP7^lo^ cells after 10 days from 3 non-carrier breast tissues. **(K)** Relative CFC number (left) and size (right) from sorted FABP7^hi^ and FABP7^lo^ cells after 10 days. Colony numbers were normalized to matched colony number from total sorted LPs (n = 8, non-carrier). P-value from unpaired student’s t-test is shown in each panel. **(L)** Representative CFC images from sorted total LPs cells 10 days after treatment with FABP7i BMS309403. **(M)** Corresponding quantification of CFC number normalized to DMSO (0 uM) control (n = 4). Significance from one-way ANOVA with Tukey post-hoc test is indicated using a letter code, with each different letter indicating p-val < 0.05. Luminal hormone sensing (HS), luminal progenitor (LP). Scale bars = 1000um.

To increase clustering resolution, we next projected these data as a graphical tree abstraction using TooManyCells (TMC), a method shown to enhance detection of unique cell states (**Figure 1B)**^16^. Briefly, in TMC, cluster resolution is data driven with a modularity cut-off based on median absolute deviations in cluster size; the leaves represent clusters with pie charts at their tips indicating the proportion of select features. TMC clustering broadly separated cells by epithelial, immune, and stromal populations, with similar representation across genotypes and minimal patient-specific branching (**Figure 1B, S1D, S1E**), except for a single patient. This sample was suspected to be in the luteal phase which we confirmed upon case review (**Figure S1F)**, underscoring menstrual cycle effects in the breast.

We next paired the CytoTRACE statistical framework^17^ with TMC to find putative progenitor populations. This predicted the luminal progenitor (LP, also known as luminal secretory^7^) subpopulation marked by lactotransferin (LTF) to be least differentiated across all epithelial lineages (**Figure 1C, S2A**). LTF expressing LP cells (LP-LTF) co-expressed many previously identified markers of LP capacity including *KIT* and *ALDH1A3* (**Figure S2B**)^18,19^. Within the LP-LTF subpopulation, leaf 80 had the highest CytoTRACE score (**Figure 1D**), predicting its greater progenitor capacity. Among the top 5 markers of LP leaf 80, fatty-acid binding protein 7 (FABP7) was almost exclusively enriched in this population (**Figure 1E, F, S2C, S2D**).

FABP7 is a fatty acid shuttle protein and a neural stem cell marker^20,21^. FABP7 preferentially binds to omega-3 derived fatty acids, and upon binding, a conformational change enables its nuclear translocation where it functions to promote nuclear acetyl-CoA production and transcription^22–24^. Pathway analysis of FABP7+ vs. FABP7- transcriptomes identified enrichment of terms related to amino acid metabolism, fatty acid synthesis and fatty acid catabolism **(Figure S2E)**. We next prepared fluorescence-activated cell sorting (FACS)-purified LP cells based on the top 5 % (FABP7^hi^) and bottom 5 % (FABP7^lo^) of FABP7-binding small molecule CDr3 signal^21^ and performed low-input proteomics^25^ on 2000 cells per fraction (**Figure 1G**, non-carrier, n=5). FABP7^hi^ cells were enriched for FABP7 at the protein level (**Figure 1H**, **Table S3**) and pathway analysis revealed lipid metabolism terms were enriched in FABP7^hi^ cells, orthogonally validating the transcriptomic analysis (**Figure 1I, S2E**). To functionally assess the progenitor potential of FABP7^hi^ LP cells, as suggested by CytoTRACE analysis, we compared the clonogenicity and colony size of FABP7^hi^, FABP7^lo^ and total LP cells in standard colony forming capacity (CFC) assays. FABP7^hi^ LP cells possessed increased clonogenic capacity and size, consistent with a progenitor cell phenotype (**Figure 1J, K**, non-carrier, n=8). Notably, addition of the FABP4 & FABP7 inhibitor BMS309403 resulted in a dose-dependent reduction in colony formation implicating a role for FABP7 in LP colony forming capacity (**Figure 1L, M,** non-carrier, n=4; FABP4 is not expressed by LPs as seen in **Figure S2D)**.

To determine whether FABP7 marks a specific cell population in the adult breast *in situ*, we used immunohistochemistry to localize FABP7 in a cohort of 31 human breast samples (**Figure 2**, n: non-carrier = 16, *BRCA1* = 7, *BRCA2* = 8). KRT14 is a ductal basal cell marker with limited expression in acinar basal cells^26^ which clearly delineated ductal basal cells for regional annotation. FABP7 cells were primarily situated within interlobular ducts, with an average of 5- 25% of interlobular ducts expressing FABP7 compared to less than 1% of acini cells across our cohort (**Figure 2A, B**). Imaging mass cytometry (IMC) was used to further co-localize FABP7 cells within a panel of key mammary cell markers^27^. We observed mutual exclusivity of FABP7 and progesterone receptor and validated the spatial localization of FABP7 cells (**Figure 2C, D**). These topographic analyses demonstrate FABP7 cells are ductal and lack luminal differentiation markers.

**Figure 2:**
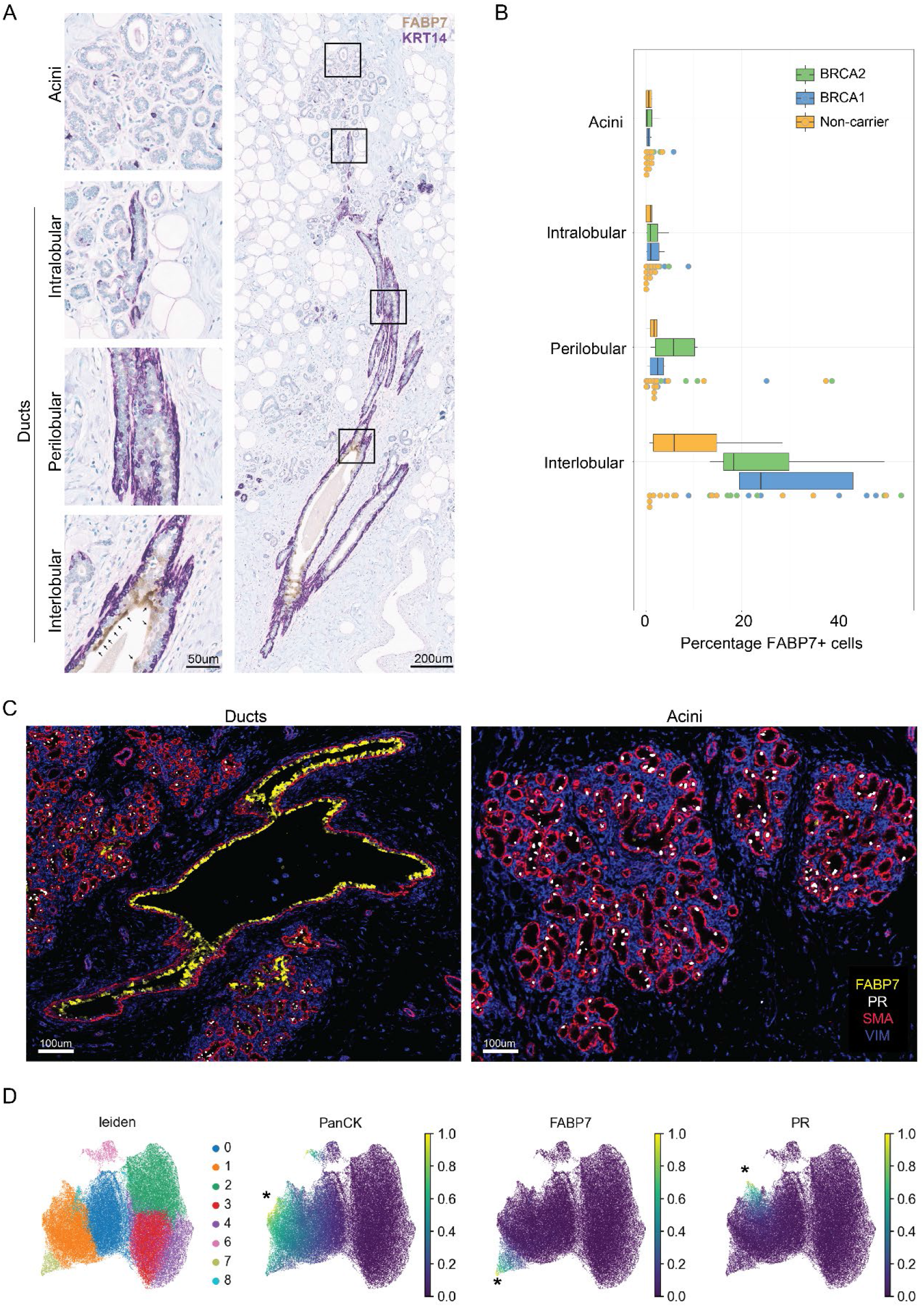
FABP7 luminal progenitors localize to the mammary ducts. (A) Representative images of histopathological annotations of human breast topographies. Acini and varying sizes of ducts were annotated (right) and FABP7 (brown, highlighted by black arrows) /KRT14 (purple) double chromogenic immunohistochemistry stained (left). **(B)** Quantification of the percentage FABP7+ cells within the total epithelium (n: non-carrier = 16, *BRCA1* = 7, *BRCA2* = 8). Mean percent positivity is shown by a black line on each boxplot with each individual data point per sample shown. **(C)** Imaging mass cytometry (IMC) detection of FABP7 (yellow), progesterone receptor (PR, luminal HS marker, white), smooth-muscle actin (basal marker, red), and stromal marker vimentin (VIM, blue). **(D)** Clustering of IMC panel signals with pan-cytokeratin+, FABP7+, and PR+ populations *highlighted.

We next investigated the dynamics of FABP7 LP cells in *BRCA1/2* mutation carriers. Our scRNAseq dataset was split into pairwise comparisons of non-carrier vs. *BRCA1* (**Figure 3**), and non-carrier vs. *BRCA2* (**Figure S3**). TMC re-clustering of *BRCA1* and non-carrier cells is shown in **Figure 3A-B**, with LP leaf 68 representing the FABP7 subpopulation within the LP-LTF lineage, which is synonymous to previous leaf 80 (**Figure 3C, D, 1F**). To infer cell state transitions, CellRank^28^ was used with the CytoTRACE kernel^29^ for unbiased rooting of the trajectory (**Figure 3E-M, O**). TMC annotations and patient features were projected onto a force-directed graph with predicted start and end macrostates (**Figure 3E-H, S4A-B**). The absorption probabilities of transitioning from the predicted start point (FABP7^hi^ LP leaf 68) to either of the 3 endpoints is shown (**Figure 3I**). This analysis predicted that unlike non-carriers, FABP7^hi^ LP leaf 68 cells transition into LP leaf 101 or LP leaf 102 in *BRCA1* carriers (**Figure 3I, J,** X^2^ = 185.61, df = 2, p-value <2.2^e-16^). LP leaf 101 and 102 cells express the LP-KRT23 cell signature yet cluster with basal cells (**Figure 3C**), resembling a mixed basal-luminal (BL) population that has lost lineage fidelity. BL cells were recently identified as a phenotype in the aging breast, which is accelerated in *BRCA* carriers^1,30^. The expression profile of LP Leaf 101/102 was consistent with previously identified markers of BL cells such as *KRT17*, *CAV1*, *KRT5*, *KRT14*, *KRT8*, *KRT18* (**Figure 3K**)^1,2,30^. Three of the top 6 genes predicted to drive the transition of LP leaf 68 ◊ 101 are known basal lineage markers including *ACTA2* and *MYLK* (**Figure 3L**). Among the top 6 predicted driver genes of the LP leaf 68 → 102 transition was *SNHG12,* a long non-coding RNA implicated in breast cancer growth and proliferation^31^ (**Figure 3M**). Further, in support of the computationally inferred trajectory of FABP7 → BL cells, immunohistochemistry revealed that K14+ BL cells were directly adjacent to FABP7+ regions in mammary ducts *in situ* (**Figure 3N)**, consistent with previous reports of BL cells in ducts^1^. This adds spatial connectivity to our *in silico* results and reiterates the absence of FABP7 in BL cells. To independently probe the FABP7 → BL cell transition in *BRCA1* carriers, we used the human breast epithelial dataset from Reed and colleagues^32^; FABP7^hi^ LP1 cluster was validated as the predicted root with a transition to BL-like LP2 cluster in *BRCA1* carriers but not in non-carriers (**Figure S5A-J**). Notably, analysis of CytoTRACE rooted trajectories in our *BRCA2* data did not yield the same predicted FABP7 → BL cell path, suggesting this transition is *BRCA1*-specific (**Figure S3A-N**). Together, FABP7 LPs are a subset of the LP-LTF lineage and are a predicted root progenitor population in the *BRCA1* high-risk breast (trajectory summarized in **Figure 3O**).

**Figure 3:**
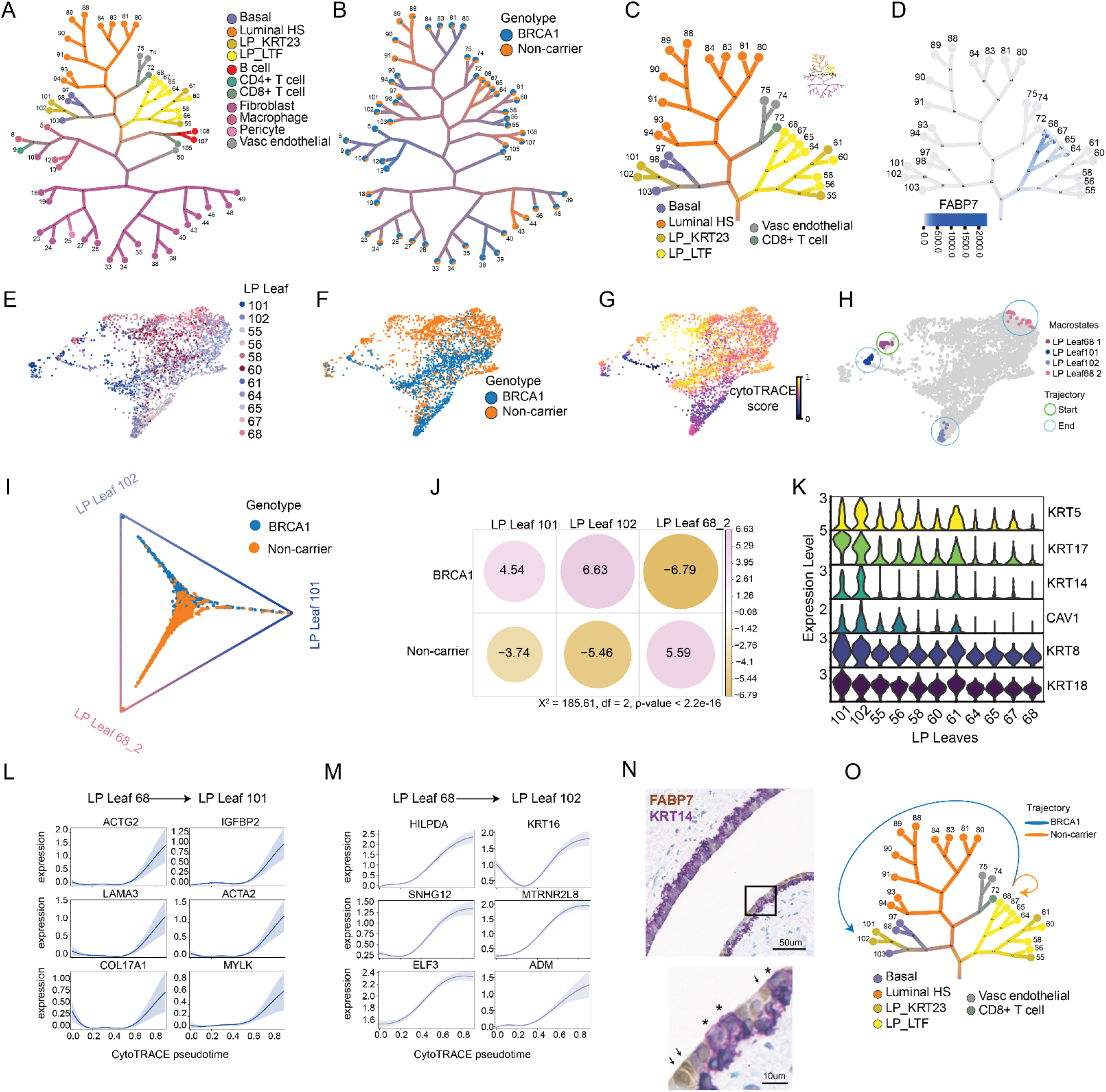
FABP7 luminal progenitors are the root of aberrant differentiation in *BRCA1* carriers. (A) TMC clustering annotated by broad cell type and **(B)** genotype for pairwise comparison of non-carrier and *BRCA1* cells. Leaf (cluster) number is shown. **(C)** TMC tree subsetted to epithelial branches and **(D)** annotated for FABP7 expression. LP Leaf 68 represents the FABP7^hi^ branch. **(E)** LP TMC annotations projected onto a force-directed graph (FDG). **(F)** FDG annotated by genotype, **(G)** CytoTRACE score, and **(H)** macrostates with start and end points denoted by open circles. **(I)** Absorption probability of transitioning into either of the three end macrostates, colored by genotype. **(J)** Dotplot of Pearson residuals from Chi-squared analysis of positive log odds towards each endpoint. **(K)** Violin plot of mixed basal-luminal marker expression across LP leaves. **(L)** Top 6 driver genes for the transition from LP Leaf 68 to LP Leaf 101. **(M)** Top 6 driver genes for the transition from LP Leaf 68 to LP Leaf 102. **(N)** Representative immunohistochemical detection of FABP7 (brown, black arrows) and KRT14 (purple) from a duct of a *BRCA1* carrier. Black asterisks mark K14+ mixed-basal luminal cells. **(O)** Summary of LP Leaf 68 trajectory in *BRCA1* (blue) or non-carrier (orange). Luminal hormone sensing (HS), luminal progenitor (LP).

BRCA1 is commonly thought of as a DNA damage repair protein yet plays a less appreciated role as a key regulator of fatty acid synthesis^33,34^. BRCA1 binds to the inactive form of acetyl-CoA carboxylase, which prevents its dephosphorylation and inhibits the conversion of acetyl-CoA to malonyl-CoA, blunting fatty acid synthesis through fatty acid synthase (FASN)^33^. Due to convergence of FABP7 and BRCA1 on regulation of acetyl-CoA pools^22,33^, we interrogated our scRNAseq data using the Compass algorithm *in silico*^35^ and predicted shifts in cellular metabolic states of FABP7 cells between non-carrier and *BRCA1* carriers. Specifically, FABP7 cells were predicted to have enhanced fatty acid metabolism in *BRCA1* but not *BRCA2* carriers when compared to non-carriers (**Figure 4A, B, S6**). Key players in fatty acid elongation, synthesis, and acetyl-CoA metabolism (*FASN*, *ACSL1*, ACAT2, and *ACLY*) were transcriptionally enriched in FABP7 cells from *BRCA1* carriers (**Figure 4C**). Given the role of fatty acid signaling in stemness in other systems^36,37^, we next tested whetherperturbation of fatty acid metabolism affects the progenitor capacity of FABP7 cells. We first treated FACS-purified total basal and total luminal progenitors with the FDA-approved FASN inhibitor TVB-2640^38^. Lineage specificity was observed with FASNi, as colony-forming capacity was only ablated in *BRCA1* LP cells at the highest dose while basal colonies remained unaffected irrespective of *BRCA* status (**Figure S7A- B**). Importantly, FACS-purified FABP7^hi^ cells displayed exquisite sensitivity to TVB-2640 in both non-carriers and *BRCA1* carriers, with a significantly greater reduction in colony-forming capacity in *BRCA1* carriers (**Figure 4D, E,** p.adj = 0.0452). This highlights the importance of fatty acid metabolism in the FABP7 LP cell population, especially in the *BRCA1* breast.

**Figure 4:**
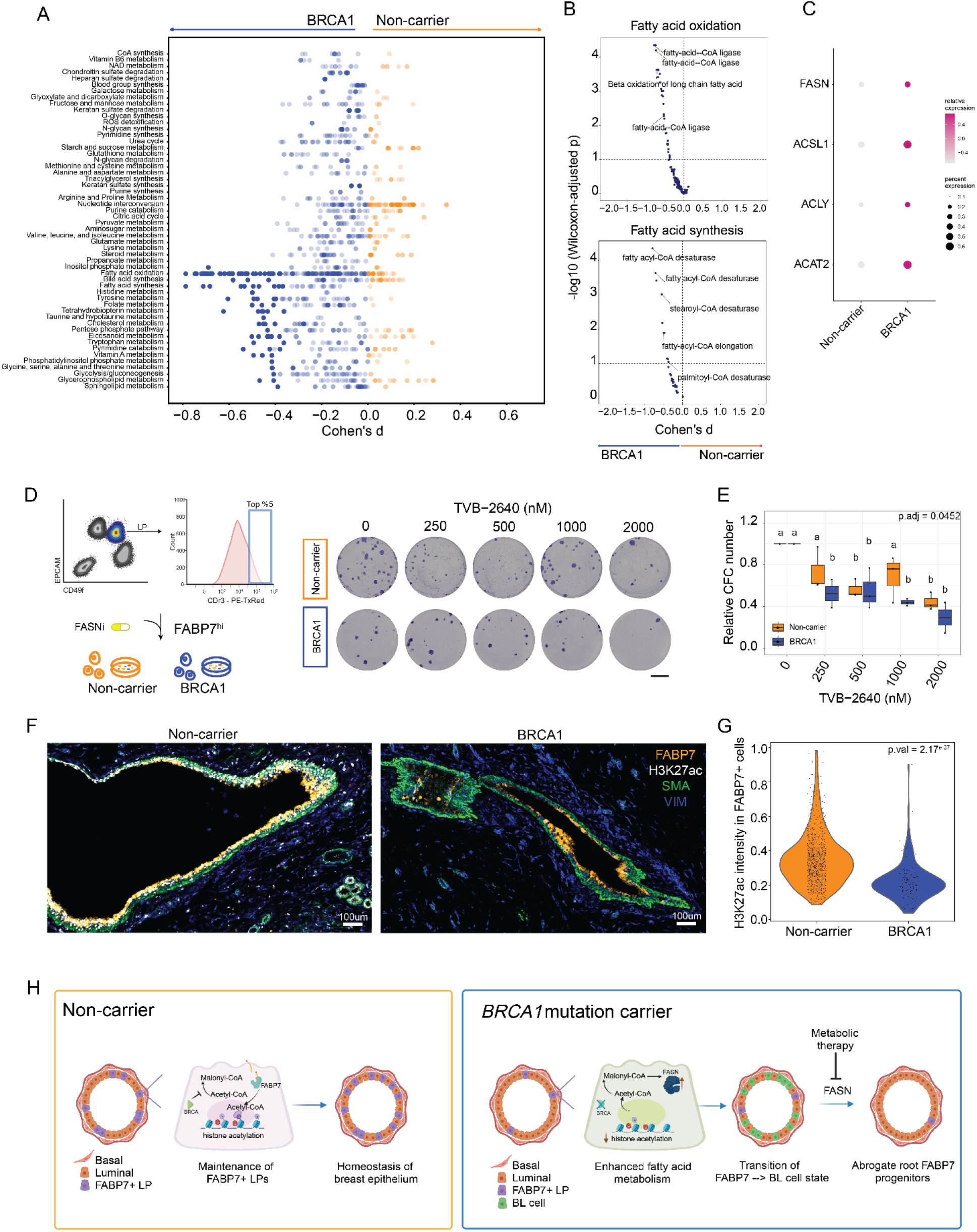
Selective metabolic vulnerability of FABP7 luminal progenitors. (A) Metabolic subsystems in non-carriers vs. *BRCA1* FABP7 luminal progenitors (LPs, n= 294 cells per group). **(B)** Specific reactions within the fatty acid oxidation (upper) and fatty acid synthesis (lower) subsystems from panel A in non-carriers vs. *BRCA1* FABP7 LPs. **(C)** Key metabolic genes enriched in *BRCA1* FABP7 cells. **(D)** Sorting schematic for isolation of FABP7^hi^ (CDr3 top 5 %) cells from *BRCA1* and non-carriers (left), and representative CFC images at day 9 post-treatment with FASNi (TVB-2640). **(E)** Colony number relative to DMSO (0 nM) control 9 days after TVB-2640 treatment (n = 3 per genotype). Two-way ANOVA with Tukey post-hoc test was performed with significance indicated by adjusted p-value for the interaction effect between genotype and a letter code indicating significance between conditions is shown. Scale bars = 1000 um. **(F)** Representative images from imaging mass cytometry (IMC) detection of FABP7 (orange), H3K27ac (white), smooth-muscle actin (basal marker, green), and stromal marker vimentin (VIM, blue). **(G)** Quantification of H3K27ac signal intensity within FABP7 positive cells between non- carrier (n=3) and *BRCA1* (n=4). Student’s unpaired t-test p-value is inset. **(H)** A model of the proposed effect of *BRCA1* monoallelic loss in FABP7 cells. The metabolic role of *BRCA1* in lipid metabolism is central to the maintenance of FABP7 progenitors. With reduced *BRCA1*, acetyl-CoA pools are dysregulated concomitant with FASN increase, which drives the transition of FABP7 LPs to a mixed basal-luminal (BL) cell state. FABP7 progenitor capacity is abrogated by metabolic perturbation through FASN inhibition, offering an avenue for therapeutic interception.

Finally, acetyl-CoA is central to cellular metabolism and a critical mediator of cell state shifts as a substrate for glycolysis, fatty-acid synthesis, and histone acetylation^39,40^. Enhanced fatty-acid synthesis with *BRCA1* heterozygosity may deplete cytoplasmic and nuclear acetyl-CoA reservoirs blunting histone acetylation, and ultimately result in the loss of lineage fidelity of FABP7 cells (**Figure 3**). To assess histone acetylation in FABP7 cells, IMC was used to co- localize H3K27ac and FABP7 within a validated breast panel^27^ (**Figure 4F, G**). Quantification of H3K27ac within FABP7 cells revealed depletion of global H3K27ac in *BRCA1* carriers (**Figure 4G**, p-value = 2.17^e-37^), consistent with the concept that histone modifications may drive the aberrant FABP7 cell state trajectories observed in the *BRCA1* high-risk breast, as modeled in Figure 4H.

## Discussion

The breast biology field has invested greatly in dissecting human breast heterogeneity with over 1 million single cells across 200 human breast tissues profiled by scRNAseq to date^2,4,7,8,32^. These compendiums reveal vast heterogeneity in the human breast identifying numerous novel epithelial and stromal cell states. Building on these studies, we find a unique FABP7 progenitor subpopulation within the LP-LTF lineage^4,7^ that are the predicted root of aberrant epithelial differentiation with *BRCA1* heterozygosity, and demonstrate that FABP7 enriches luminal progenitor capacity. We pinpoint this population in mammary ducts *in situ*, and FABP7 enables the purification of a ductal luminal progenitor subset for the first time. The ductal distribution pattern of FABP7 *in situ* is similar to GPR125+ basal progenitors and luminal label retaining cells that were reported in mice^41,42^, supporting breast ducts as an untapped reservoir for adult progenitors in mouse and human.

FABP7 cell proteomes, transcriptomes, and functional assays revealed these cells are highly equipped with fatty acid machinery that is paramount to their progenitor capacity, as they show vulnerability to FASN inhibition. FASN was previously suggested as a mediator of *BRCA1*- associated metabolic syndrome^33,34^, and FABP7 progenitors may mechanistically be a cellular target for FASNi in this context. Epidemiological studies have correlated dietary intake of omega- 3 fatty acids from fish, canola, and linseed oils with reduced breast cancer incidence in pre- and post-menopausal women, with experimental murine models reiterating similar effects^43–46^. In addition to FASNi, modulating dietary omega-3 intake may be worth investigating in the epigenetic regulation of the *BRCA1* breast. In neural systems, FABP7 localized to the nucleus upon binding omega-3^47^, mediating nuclear acetyl-CoA production through ACLY to maintain histone acetylation^22^. The observed reduction in global histone acetylation within ductal FABP7 cells from *BRCA1* carriers centers the BRCA1-acetyl-CoA axis as pertinent to mammary cell dysregulation in the high-risk breast. Altogether, this study uncovers a unique FABP7 progenitor population in the human breast that is spatially restricted, dysregulated in *BRCA1* mutation carriers, and responds to FASNi, providing a new targetable epithelial population for breast cancer chemoprevention strategies in high-risk women.

## Methods

### Patient samples

All human tissues were acquired with patient consent and approval from the Research Ethics Board of University Health Network (Toronto, ON). Menstrual stage (premenopausal, follicular and luteal) was determined by histological examination of breast specimen stained with hematoxylin & eosin from at minimum 5 tissue blocks per patient^48^. Staging was confirmed by pathologist (H. Berman). Breast specimens were collected from true prophylactic mastectomy or contralateral unaffected breast tissues, with necessary patient information presented in **Table S1**. Breast specimens were minced and enzymatically dissociated in DMEM:F12 1:1 media with 15 mM HEPES plus 2 % BSA, 1 % penicillin-streptomycin, 5 ug/ml insulin, 300 U/ml collagenase (Sigma, C9891) and 100 U/ml hyaluronidase (Sigma, H3506) shaking gently at 37 °C for 16-18 hours. Epithelial, endothelial, and fibroblast-rich fractions were harvested by centrifugation at 80 g (30 seconds), 200 g (4min), and 450 g (5min), respectively, and then viably cryopreserved, as described previously^11,49^.

### Human breast single cell suspensions for FACS and MULTI-seq

Viably cryopreserved human breast tissues were thawed and dissociated into single cell suspensions using standard methodology for human breast tissue^11^. Briefly, cell clusters were triturated in 0.25% trypsin-EDTA (Stem Cell Technologies, 07901) followed by 5 U/ml dispase (Stem Cell Technologies, 07913) and 50 ug/ml DNase I (Sigma, D4513). Cells were then washed with HBBS + 2% FBS and filtered using a 40 um strainer. For MULTI-seq, a second 40 um filtering step was performed prior to sample barcoding as described below.

### Human breast sample multiplexing and MULTI-seq library prep

Samples were multiplexed using lipid-tagged indices with established protocols^50^. Briefly, a single-cell suspension of each breast fraction (epithelial, endothelial, and fibroblast-rich) were generated as described above, and then washed twice with 1x PBS (Wisent Bio, 311-010-CL). 1.5x10^5^ cells per fraction were pooled per patient prior to MULTI-seq labelling (n = 4 patients per batch). Each pooled sample was given a unique MULTI-seq oligonucleotide barcode following the standard MULTI-seq workflow. Excess oligonucleotide was quenched with 1 mL 1 % BSA in 1x PBS solution. Each sample was spun down at 400g for 5min and supernatant removed. A second wash was performed similarly, and cells were suspended in a final solution of 0.5 % BSA in 1x PBS. Each sample was counted and then pooled at a concentration necessary to target 2x10^4^ cells in the final transcript library, and standard protocols for the 10x Genomic Single Cell 3’ RNA kit (v3 chemistry) were used. Sequencing was performed using the NovaSeq 6000 (Illumina) as per the standard 10x configuration.

### FACS purification of FABP7^hi^ cells

Human breast epithelial single cell suspensions (as above) were incubated with 1 uM CDr3 in DMEM:F12 at 37 °C for 1 h, followed by a 1 h de-stain in DMEM:F12 at 37 °C. Cells were then stained with standard antibodies for immune and vascular lineage exclusion: anti-CD45-PE/Cy7 (BioLegend, 304016, 1:200), anti-CD31-PE/Cy7 (BioLegend, 303118, 1:50), and for mammary epithelial cells subsets: anti-CD326 (EpCAM)-PE (BioLegend, 324206, 1:50) and anti-CD49f-FITC (BioLegend, 313606, 1:100). Live cells were selected based on Dapi (Sigma, D9542, 1:10000) exclusion. Samples were sorted into PBS for proteomics or 1x HBSS + 2% FBS for colony-forming assays. FACS was performed using the BD™ FACSAria Fusion.

### Proteomic sample processing

Cells were deposited into wells of Teflon-coated slides (Tekdon, template available upon request) by pipette or directly by FACS. The solution accompanying the cells was allowed to evaporate at room temperature in a biosafety cabinet and then slides were stored at -80 °C until further processing. For lysis and protein reduction, 2.5 μL of buffer consisting of 30% (v/v) Invitrosol (ThermoFisher, MS10007), 15% (v/v) acetonitrile, 0.06% (w/v) n-dodecyl-β-D-maltoside (Sigma, 850520P), 5 mM Tris(2-carboxyethyl)phosphine, and 100 mM ammonium bicarbonate in HPLC grade water. Slides were placed in a 37 °C humidity chamber and allowed to incubate for 30 min. For alkylation and protein digestion, 1 μL of buffer containing 35 mM iodoacetamide and Trypsin/Lys-C (Promega, V5072) was added to each sample. Samples were allowed to digest for 2 h in a 37 °C humidity chamber. Digestion was quenched by bringing samples to 0.1% (v/v) formic acid with 8 μL of 0.14% (v/v) formic acid in 37 °C HPLC grade water and then samples were transferred to 96-well plate. The wells were washed with an additional 8 μL of 0.1% (v/v) formic acid and then the plate was transferred to EASY-nLC™ 1000 System chilled to 7 °C.

### Mass spectrometry data acquisition

LC-MS/MS analysis was performed on an Orbitrap Fusion MS (ThermoFisher) coupled to EASY-nLC™ 1000 System (ThermoFisher). Peptides were washed on pre-column (Acclaim™ PepMap™ 100 C18, ThermoFisher) with 60 μL of mobile phase A (0.1% FA in HPLC grade water) at 3 μL/min separated using a 50 cm EASY-Spray column (ES803, ThermoFisher) ramping mobile phase B (0.1% FA in HPLC grade acetonitrile) from 0 % to 5 % in 2 min, 5 % to 27 % in 160 min, 27 % to 60 % in 40 min interfaced online using an EASY-Spray™ source (ThermoFisher). The Orbitrap Fusion MS was operated in data dependent acquisition mode using a 2.5 s cycle at a full MS resolution of 240,000 with a full scan range of 350-1550 m/z with RF Lens at 60 %, full MS AGC at 200 %, and maximum inject time at 40 ms. MS/MS scans were recorded in the ion trap with 1.2 Th isolation window, 100 ms maximum injection time, with a scan range of 200-1400 m/z using Normal scan rate. Ions for MS/MS were selected using monoisotopic peak detection, intensity threshold of 1,000, positive charge states of 2-5, 40 s dynamic exclusion, and then fragmented using HCD with 31 % NCE.

### Mass spectrometry raw data analysis

Raw files were analyzed using FragPipe (v.20.0) using MSFragger^51,52^ (v.3.8) to search against a human (Uniprot, 43,392 sequences, accessed 2023-02- 08) proteome – canonical plus isoforms. Default settings for LFQ workflow^53,54^ were used using IonQuant^55^ (v.1.9.8) and Philosopher^56^ (v.5.0.0) with the following modifications: Precursor and fragment mass tolerance were specified at -50 to 50 ppm and 0.15 Da, respectively; parameter optimization was disabled;; MaxLFQ min ions was set to 1; MBR RT tolerance was set to 2 min, and MBR top runs was set to 100.

### Colony-forming Assays

FACS purified single cell suspensions (500 cells/per well) were seeded in Human EpiCult-B media (Stem Cell Technologies) supplemented with 5 % FBS on day 0, on a feeder layer of 20,000 irradiated NIH 3T3 cells in collagen-coated 6-well dishes (Greiner). On day 1, media was changed to FBS-free EpiCult-B media and cultures were undisturbed for 9-10 days with 5 % O2 at 37 °C. Drug treatments (FASNi TVB-2640, Cayman No. 35703 or FABP7i BMS309403 Cayman No. 10010206) were added at the same time as the media change on day 1 with a final DMSO concentration <1 % total volume per well for each drug concentration. All control wells (denoted as 0 uM in figures) were treated with the highest DMSO concentration in the dilution series to control for DMSO effects. After 9-10 days, cells were quickly fixed with 1:1 acetone:methanol, air dried, and Giemsa stained. Colonies were visualized using the Biotek Cytation 5 with Gen5 software (3.14). Colonies for manually annotated in QuPath (0.3.0) software. For the comparison of FABP7^hi^ vs. FABP7^low^ colony forming capacity, colony counts were normalized to the total colonies formed by total sorted LP cells per patient. Drug-treated colonies were normalized to the matched DMSO-only (denoted as 0 uM) control. One-way ANOVA (FABP7i) or Two-way ANOVA with Tukey post-test (FASNi) was performed to test either the effect of the drug alone or for interaction effects between lineage or genotype in addition to the drug effect, respectively. Significance is displayed using a letter code generated from the multcompView R package (v. 0.1.9).

### Immunohistochemistry (IHC) and QuPath

Histopathologic assessment of human breast tissues was first performed using hematoxylin & eosin to confirm no malignancy was present and for menstrual cycle staging. All IHC experiments were done on 4 um sections of formalin-fixed paraffin embedded tissues. Sections were deparaffinized and dehydrated in a xylene and alcohol gradient, followed by pressurized antigen retrieval for 8 min (Reveal Decloaker, pH 6.0). Endogenous peroxidases were blocked by incubation with 3 % H2O2 and then blocked for 1 h (Dako Serum-free protein block). Sections were then incubated with the 1^st^ primary antibody (FABP7, Abcam No. ab279649, 1:700) overnight at 4 °C. After incubation with appropriate secondary HRP for 1 h at room temperature, FABP7 was developed using diaminobenzidine for 5 min. HRP was then quenched using Bloxall (Dako) and incubated with 2^nd^ primary antibody (KRT14, Abcam No. ab233910, 1:500) for 1 h at room temperature. After incubation with appropriate secondary HRP, KRT14 was developed using ImmPact VIP Purple (Vector, VECTSK4605) for 10 min. Images were captured using Aperio Scanscope. Ducts and acini were manually annotated on each slide using QuPath software, guided by a breast pathologist and aided by KRT14 staining. FABP7 localization was assessed by building a custom classifier. The custom classifier is a composite classifier built from two separate trained classifiers. The first classifier was built by setting stain vectors to specifically identify KRT14+ cells (purple) from KRT14- cells. The second classifier separates identified KRT14- cells into either FAPB7+ (brown) and FAPB7- cells based on signal intensity as determined by selected stain detection thresholds, ultimately providing a proportion of FAPB7+ cells per epithelial annotation.

### Imaging Mass Cytometry Panel

An 11-marker panel (**Table S4**) was designed to target cell types known to exist in breast tissue, including markers of luminal (CK8/18, PR, PanCK, E-Cadherin, FABP7), basal (SMA, PanCK, E-Cadherin), and stroma (Vimentin, CD31, COL1A2) to outline overall tissue architecture. Proliferation (Ki67) and histone acetylation (H3K27ac) markers were also included. Each antibody was conjugated to a lanthanide-series metal isotope of a unique mass to enable detection using IMC. Conjugations were performed using the MaxPar antibody labeling kits (Standard Biotools) according to MaxPar labeling protocol.

### Imaging Mass Cytometry preparation and acquisition

Formalin-fixed and paraffin-embedded tissue samples were baked at 60 °C degrees for 1 hour to enhance adherence of the tissue to the slide. Slides were deparaffinized in 3 x 10 min solutions of xylene and rehydrated in a graded series of alcohol (ethanol: deionized water 100:0 (x2), 96:4, 90:10, 80:20, 70:30, 5 min each). Heat-induced epitope retrieval was performed using a Decloaking ChamberTM NxGen (Biocare Medical) for 30 min at 90°C in HIER buffer (10mM Tris Base, 1mM EDTA, pH 9.2). Slides were allowed to cool at room temperature and then blocked using blocking buffer (3% BSA and 5% horse serum in TBS) for 1 h. Samples were incubated overnight at 4 °C in unconjugated primary FABP7 and PR antibodies diluted in blocking buffer. Tissue samples were washed 3 x 5 min in TBS to remove unbound primary antibody and incubated in metal conjugated anti-mouse and anti-rabbit secondary stains diluted in blocking buffer for 1 h at room temperature. Samples were once again washed 3 x 5 min in TBS to remove unbound secondary antibody and incubated overnight at 4 °C in primary metal-conjugated antibody cocktail diluted in blocking buffer. Tissue samples were stained using iridium (a DNA intercalator, at 1:1000 for 5 min) and washed 3 x 5 min in TBS before being dried for IMC acquisition. Images were acquired using an XTi imaging system (Standard Biotools). The tissue regions of interest consisting of either breast ducts or alveoli were selected and laser-ablated in a 1 um rastered pattern at 800 Hz, and preprocessing of the raw data was completed using commercial acquisition software (Standard Biotools).

### Imaging Mass Cytometry data analysis

Data were converted to TIFF format before further processing. Individual cells were segmented using Mesmer and a mask was created for each segmented image. Single-cell segmentation masks and TIFF images for each of the 11 channels were overlaid and the mean expression levels of markers and spatial features of single cells were quantified. This resulted in a measurement file storing the single cell expression of every marker panel summarized by mean pixel intensity. Prior to analysis, the single cell data were censored at the 99th percentile to remove outliers, and then scales normalized by dividing each channel by the channel’s maximum value. No further transformations were performed. The Scanpy (v1.9.5) package was used for data visualization. For UMAP creation, Leiden clustering and z-scores were used to visualize marker expressions in distinct clusters. To visualize the difference in the acetylation of FABP7+ cells, gating was performed to retain only cells with FABP7 expression of >0. The distribution of H3K27ac in these cells was then visualized in a violin plot via the Scanpy package to display differences between non-carrier and *BRCA1* carrier samples.

### Bioinformatics Analysis of MULTI-seq data

#### MULTI-seq data processing

Raw sequencing reads were aligned using CellRanger 3.1.0 with the GRCh38-3.0.0 human genome as reference. MULTI-seq data quality-control was performed using the R packages deMULTIplex (v1.0.2)^50^, DoubletFinder (v3)^57^, and Seurat (v4.3.0)^58^. A combination of demultiplex and DoubletFinder was used to collectively remove homotypic and heterotypic doublets from downstream analysis. A mitochondrial gene expression cutoff of 15% was used, with >200 unique genes per cell. Unsupervised clustering (resolution 0.8) identified eleven broad clusters that were first visualized using Harmony (v.0.1.1). Post-QC raw counts matrix was then used as input to TooManyCells (v.2.0.0.0) with default thresholds of min cell 250 and min features 1 for hierarchical spectral clustering. The first instance of TooManyCells was run on a high-performance cloud computing cluster and downstream analysis performed using TooManyCells (v.2.0.0) in Ubuntu (20.04) within a Windows operating system. The resulting tree was then pruned using 4 median absolute deviations of leaf size. These same cutoffs were used for all instances of TooManyCells visualization (all samples, non-carrier vs *BRCA1*, and non-carrier vs. *BRCA2*). The leaf annotation numbers were then imported into the matched Seurat object and cell types were assigned using the ScType R package (https://github.com/IanevskiAleksandr/sc-type)^59^ with a curated marker set (**Table S2**). TMC leaf annotations used for cluster identity in ScType analysis.

#### Progenitor prediction

The CytoTRACE computational framework^17^ was used to infer progenitor cell states. CytoTRACE scores were calculated for all epithelial cells (broadly basal, luminal progenitor, and luminal mature), with a high CytoTRACE score predicting a cell population with higher progenitor features. Batch was corrected for using ComBat for CytoTRACE analysis. TMC leaf marker genes were generated using Seurat FindAllMarkers function with logFC >0.25 and p<0.05 with expression in a minimum of 5% of cells.

#### Single Cell Pathway Analysis

Pathway analysis was performed using SCPA (1.2.0) based on multivariate distribution of Reactome pathways across groups (msigdbr v.7.5.1), which assigns a q-value that represents differential activity^60^. A q-value cutoff of 2.5 with adjusted p-value <0.01 used for selecting pathways for visualization with fold change presented between FABP7+ vs - cells. FABP7+ cells were annotated based on having any detectable counts for FABP7.

#### Cell state transition inference

Prediction of cell state transitions across genotypes was performed using the CellRank python package (v1) using the CytoTRACEKernal for unbiased root selection^28^. TMC leaf annotations were imported and projected onto a force-directed graph. Genes were filtered based on minimum detection in 10 cells. Gene expression was then imputed to calculate differentiation potential. A transition matrix was computed using the ‘hard’ threshold scheme that is based on the Palantir algorithm^61^. Shur decomposition (20 components) with the Generalized Perron Cluster Cluster Analysis (GPCCA) estimator was used to compute macrostates. The number of macrostates were selected after visualizing the real part of the top 20 eigenvalues of the real part and selecting the number of states prior to a prominent eigengap with a minimum of 4 states (**Figure S3A** – 4 macrostates, **S4E** – 4 macrostates, **S5I** – 5 macrostates). Start and end points of the trajectory were selected by the stationary distance of the coarse-grained transition matrix, with macrostates having a lower stationary distance assigned as a start point. Absorption probabilities of transitioning to assigned end states were calculated and visualized using a circular projection plot. To determine the significance of inferred macrostate trajectories, the log odds ratio was calculated from the absorption probability towards each end macrostate and the number of cells with positive log odds to each macrostate were tabulated to generate a 2x3 cross table and Chi-squared test was applied. The significance and Pearson residuals are presented in **Figures 3J**, **S4I**, and **S5M**. Finally, driver genes from root LP leaf 68 to LP leaf 102 were calculated using CytoTrace pseudotime as input to fit a generalized additive model (GAM). The luteal phase sample was excluded from CellRank analysis for the comparison of non-carrier vs. *BRCA2*.

#### Metabolic inference

*In Silico* single-cell flux balance analysis (FBA) was performed using Compass algorithm^35^. As Compass requires linear scaling, subsetted FABP7+/- LP cells were normalized using the Seurat NormalizeData function with relative count ‘RC’ normalization and a scale factor of 10,000. With standard settings, Compass ran for 17 h using 8 cpu cores on 294 cells per group. The resulting FBA reaction penalties for each RECON2 metabolic subsystem were negative log transformed and Wilcoxon rank-sum test was used to identify reaction subsystems predicted to be more active in each genotype. The magnitude of predicted activity is shown as a Cohen’s d value that represents the difference in mean divided by the pooled standard deviation.

### Data Availability

Processed and raw UMI count matrices, as well as raw fastq files for both barcode and transcript libraries will be deposited in the NCBI Gene Expression Omnibus. Mass spectrometry data will be available at MassIVE. Unique code will be available on Github.

## Tables

Table S1: Human breast sample summary

Table S2: Human breast cell type markers

Table S3: Differential expression of FABP7^hi^ vs FABP7^lo^ proteomes

Table S4: Imaging mass cytometry antibody panel

## Acknowledgements

We thank Drs. Federico Gaiti and Paul Waterhouse for their comments on this study. We thank Dr. Zev Gartner for providing MULTI-seq reagents. We thank Dr. Young-Tae Chang for providing the CDr3 probe. We further thank Arian Khadani from the PM flow core for their helpful contributions. We thank Dr. Mohammad Mazhab-Jafari for TVB-2640 and Dr. Farnoosh Abbas Aghababazadeh for statistics guidance. Schematics were created with BioRender.com. This work is supported by funding from Canadian Institutes of Health Research (CIHR), Breast Cancer Canada, Canadian Cancer Society Research Institute, and the Terry Fox Research Institute. T.K. and R.K. were supported through the Canadian Research Chair program. BZ and FV are supported by Ontario Graduate Scholarships. MW was supported by a CIHR postdoctoral fellowship. CWM was supported by a Banting Postdoctoral Fellowship.

## Author contributions

Conceptualization: CWM and RK

Methodology: all authors

Formal analysis: CWM, BZ, GA, MW, FV

Resources: RK, TK, HJ, HB, SH

Writing: CWM and RK

Funding acquisition: RK, TK, HJ

---

**Figure S1:**
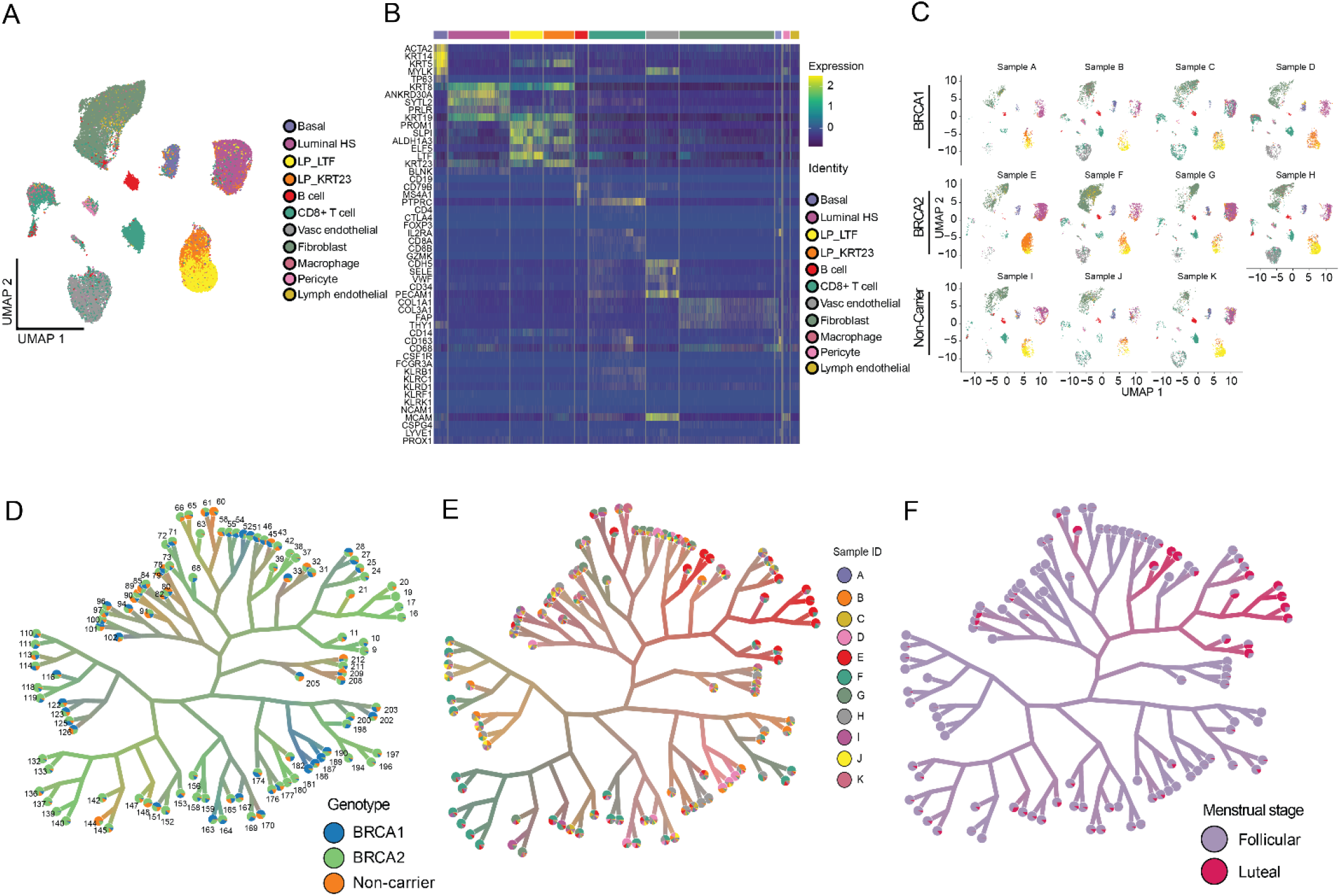
Expanded features of the MULTI-seq human breast dataset. (A) UMAP embedding of MULTI-seq dataset annotated by broad cell type. **(B)** Heatmap of cell type marker gene expression that was used as input to ScType for cell type annotation. **(C)** UMAP embedding split by patient showing representation of each cell type across all patients for *BRCA1*, *BRCA2*, and non-carriers. Clusters are annotated by cell type. TMC tree annotated by **(D)** genotype, **(E)** patient, and **(F)** menstrual stage.

**Figure S2:**
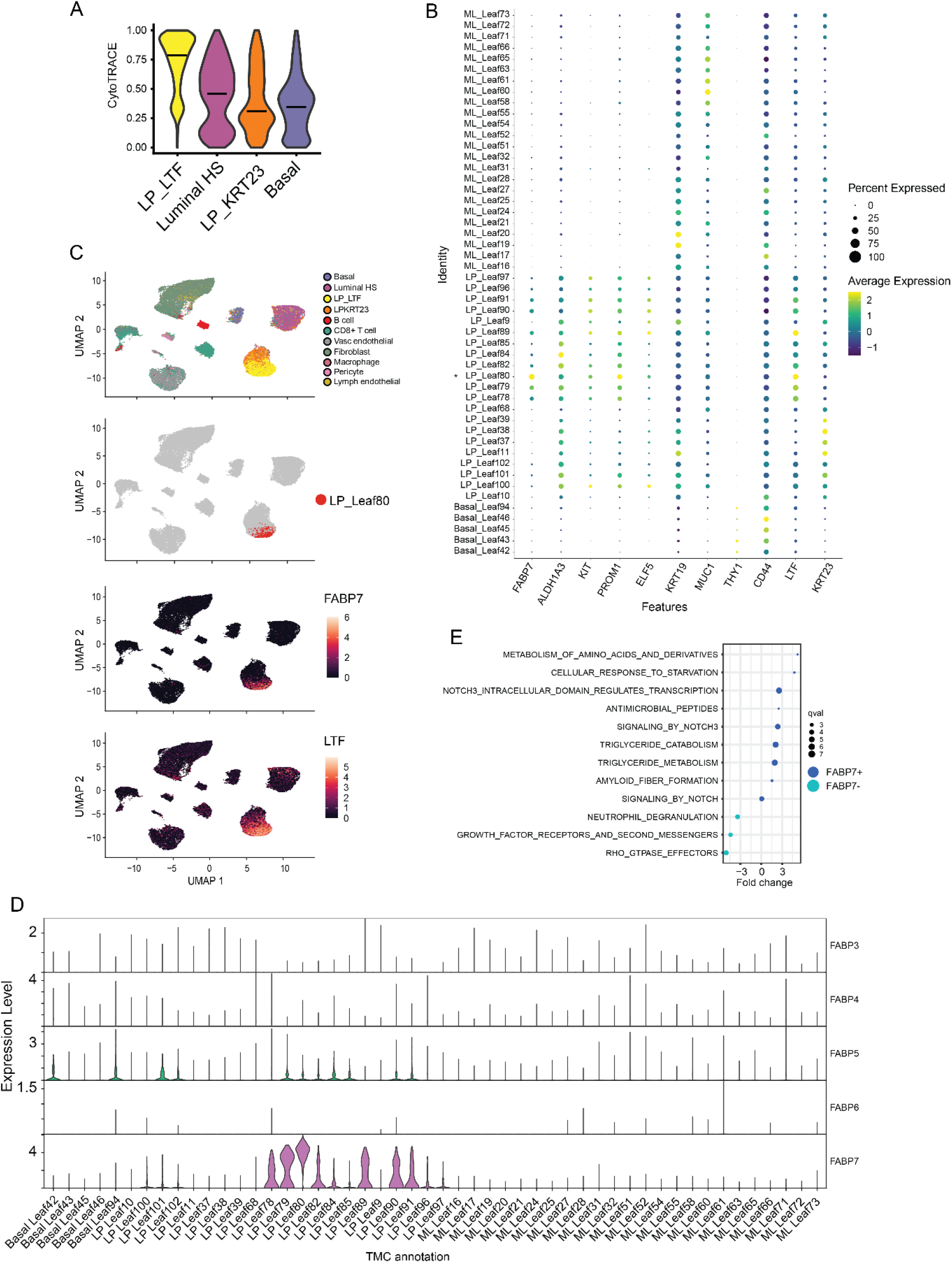
FABP7+ progenitor subset lies within LTF+ luminal progenitors. (A) Violin plot of CytoTRACE score per broad epithelial lineage. **(B)** Dot plot of previously identified human breast progenitor markers across all epithelial TMC leaves. **(C)** Localization of LP Leaf 80 cells on UMAP embeddings. FABP7 and LTF expression are also projected. **(D)** Violin plot of FABPs expression across all epithelial TMC leaves. **(E)** Pathway analysis of FABP7+ vs FABP7- LP-LTF cells. Pathways with q- val >2.5 are shown with fold change of each pathway presented. Mature Luminal (ML) = Luminal HS.

**Figure S3:**
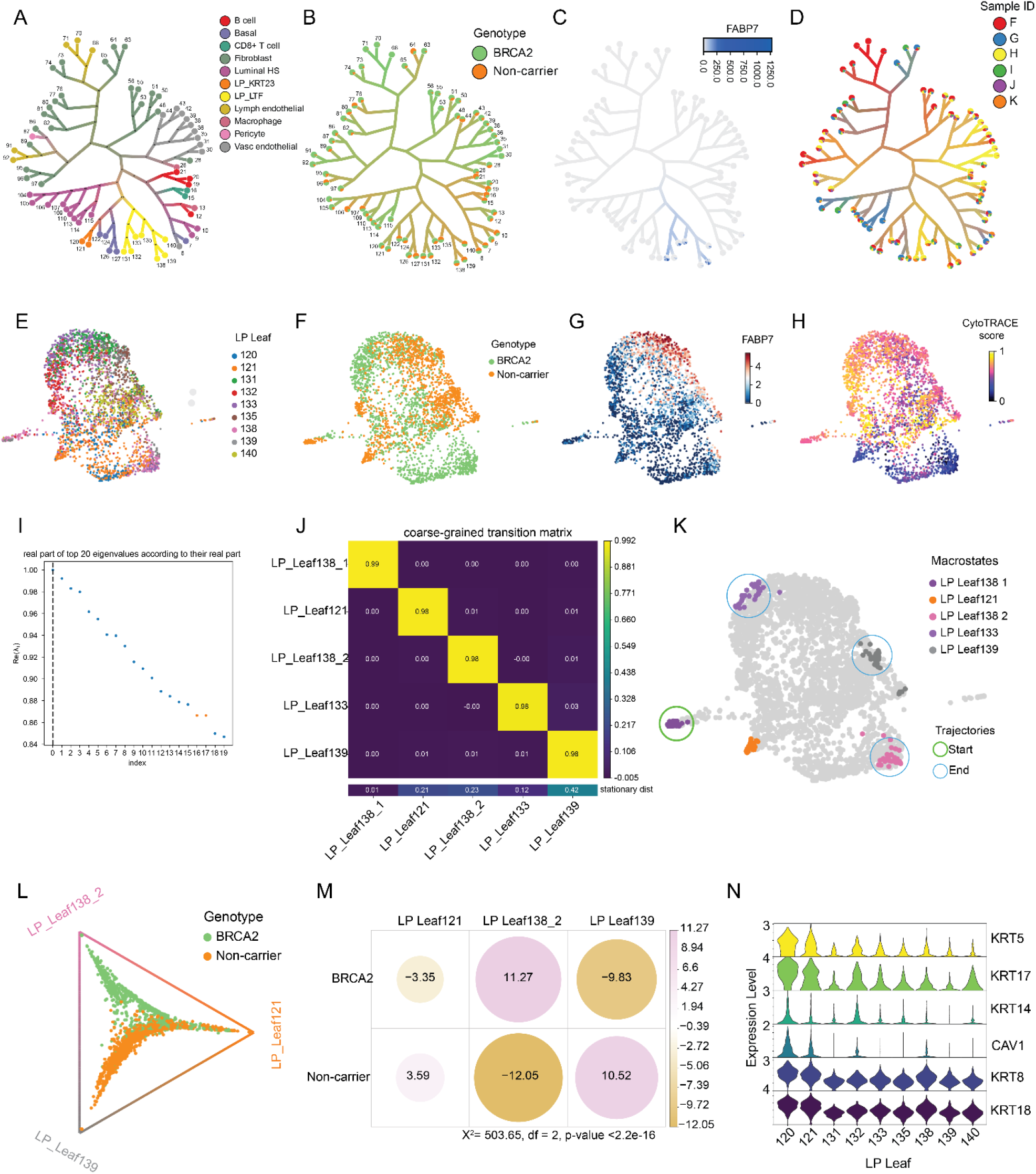
Cell state transition inference in pairwise *BRCA2* and non-carrier luminal progenitor. (**LP) cells**. **(A)** TMC clustering annotated by broad cell type and **(B)** genotype for pairwise comparison of non-carrier and *BRCA2* cells. Leaf (cluster) number is shown. TMC tree annotated for **(C)** FABP7 expression and **(D)** sample ID. **(E)** LP TMC annotations, **(F)** genotype, **(G)** FABP7 expression, and **(H)** CytoTrace score projected on force-directed graphs. **(I)** The computed real part of the top 20 eigenvalues according to their real part. Five macrostates were selected. **(J)** Coarse-grained transition matrix of CytoTRACE rooted pseudotime enabling selection of start (low stationary distance) and endpoints (high stationary distance). **(K)** Macrostates with start and end points indicated by open circles, projected onto a force-directed graph. **(L)** Absorption probability of transitioning into either of the three end macrostates, annotated by genotype. **(M)** Dot plot of Pearson residuals from Chi-squared analysis of positive log odds towards each endpoint. **(N)** Violin plot of mixed basal-luminal marker expression across LP leaves.

**Figure S4:**
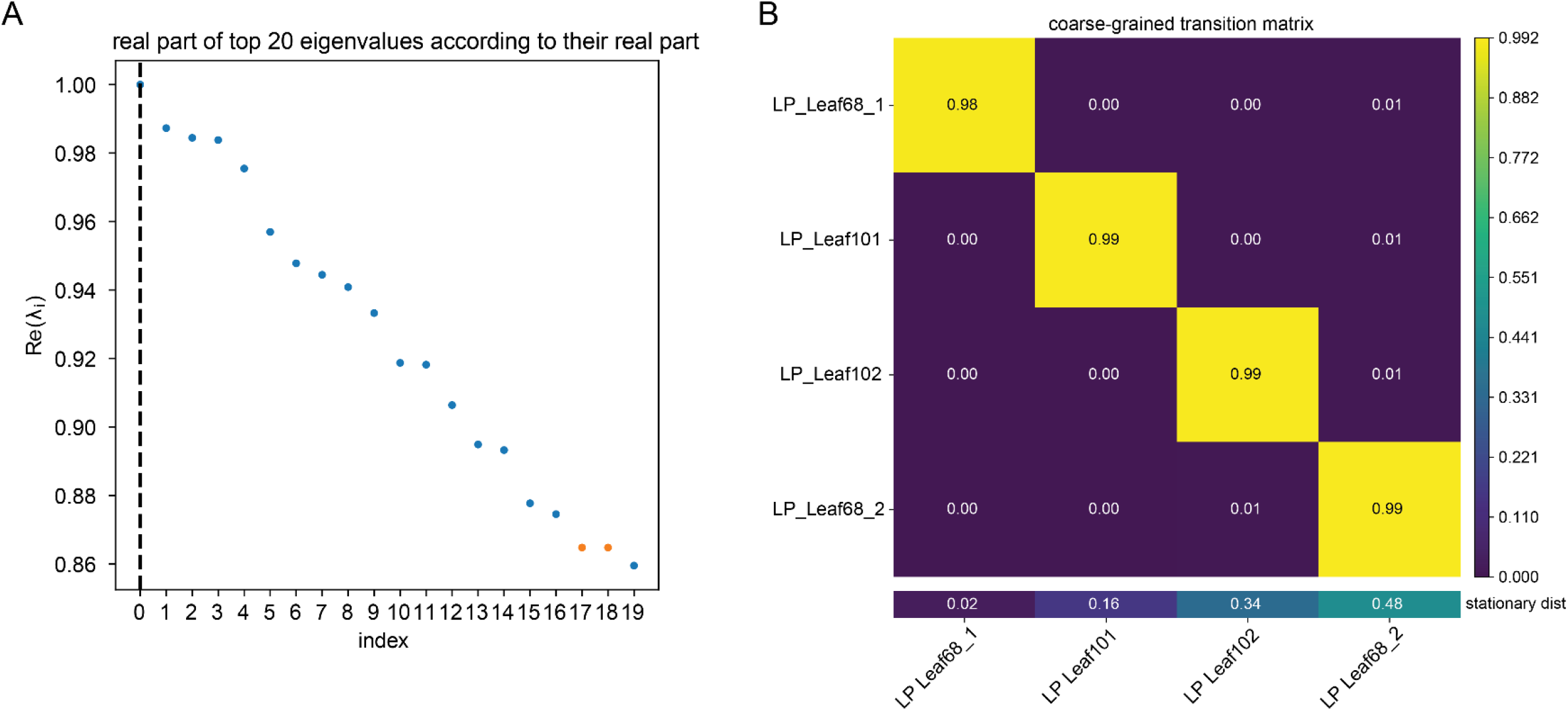
Selection of macrostates for cell state transition analysis in non-carrier and *BRCA1* carriers.(A) The computed real part of the top 20 eigenvalues according to their real part. Four macrostates were selected. **(B)** Coarse-grained transition matrix of CytoTRACE rooted pseudotime enabling selection of start (low stationary distance) and endpoints (high stationary distance).

**Figure S5:**
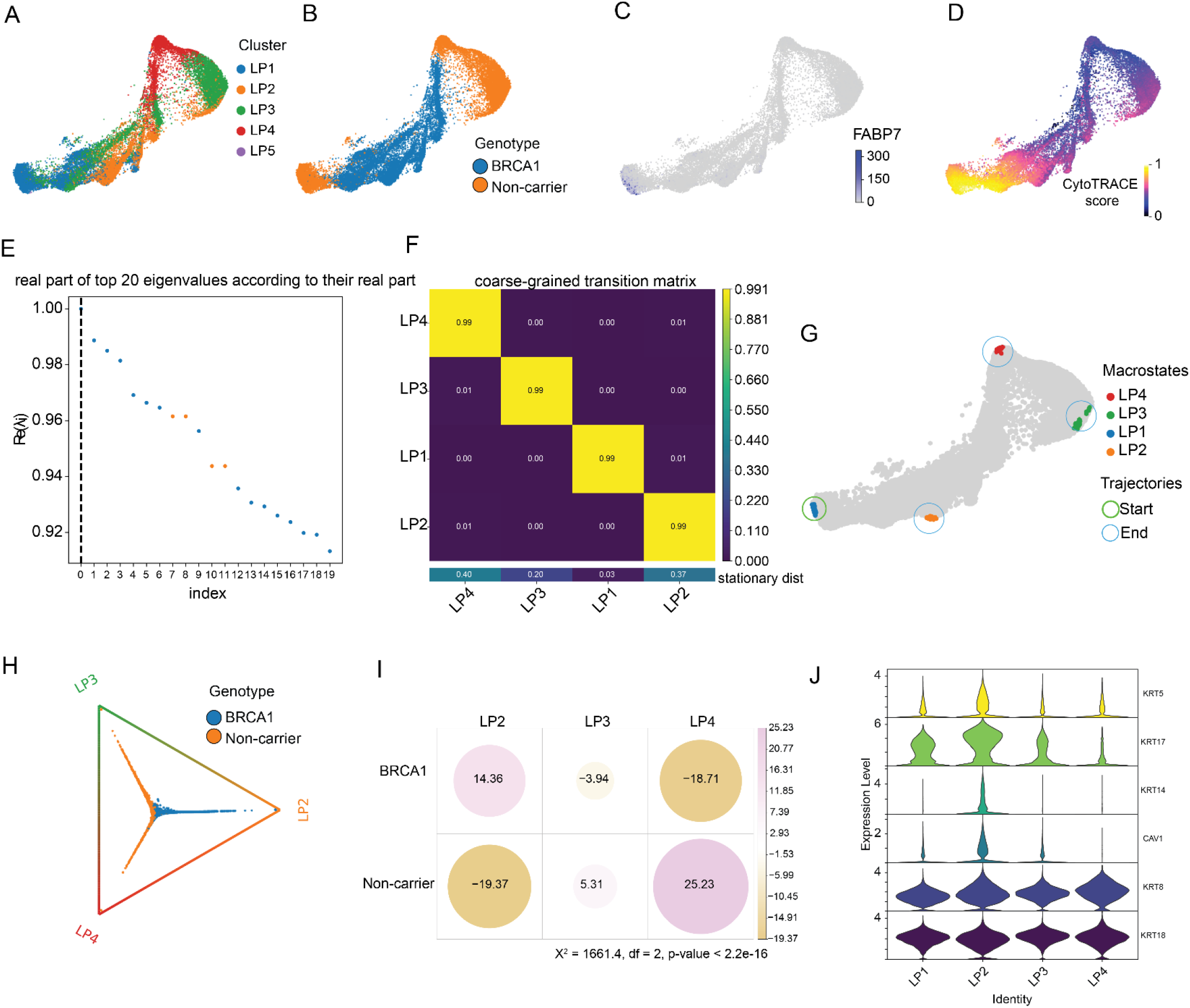
FABP7 cell state transition inference in Reed et al. 2023 human breast single-cell RNA-seq dataset. (A) Luminal progenitor (LP), **(B)** genotype, **(C)** FABP7 expression, and **(D)** CytoTRACE score projected on force directed embeddings. **(E)** The computed real part of the top 20 eigenvalues according to their real part. Four macrostates were selected. **(F)** Coarse-grained transition matrix of CytoTRACE rooted pseudotime enabling selection of start (low stationary distance) and endpoints (high stationary distance). **(G)** Macrostates with start and end points indicated by open circles, projected onto a force-directed graph**. (H)** Absorption probability of transitioning into either of the three end macrostates, annotated by genotype. **(I)** Dot plot of Pearson residuals from Chi-squared analysis of positive log odds towards each endpoint. **(J)** Violin plot of mixed basal-luminal marker expression across LP clusters.

**Figure S6:**
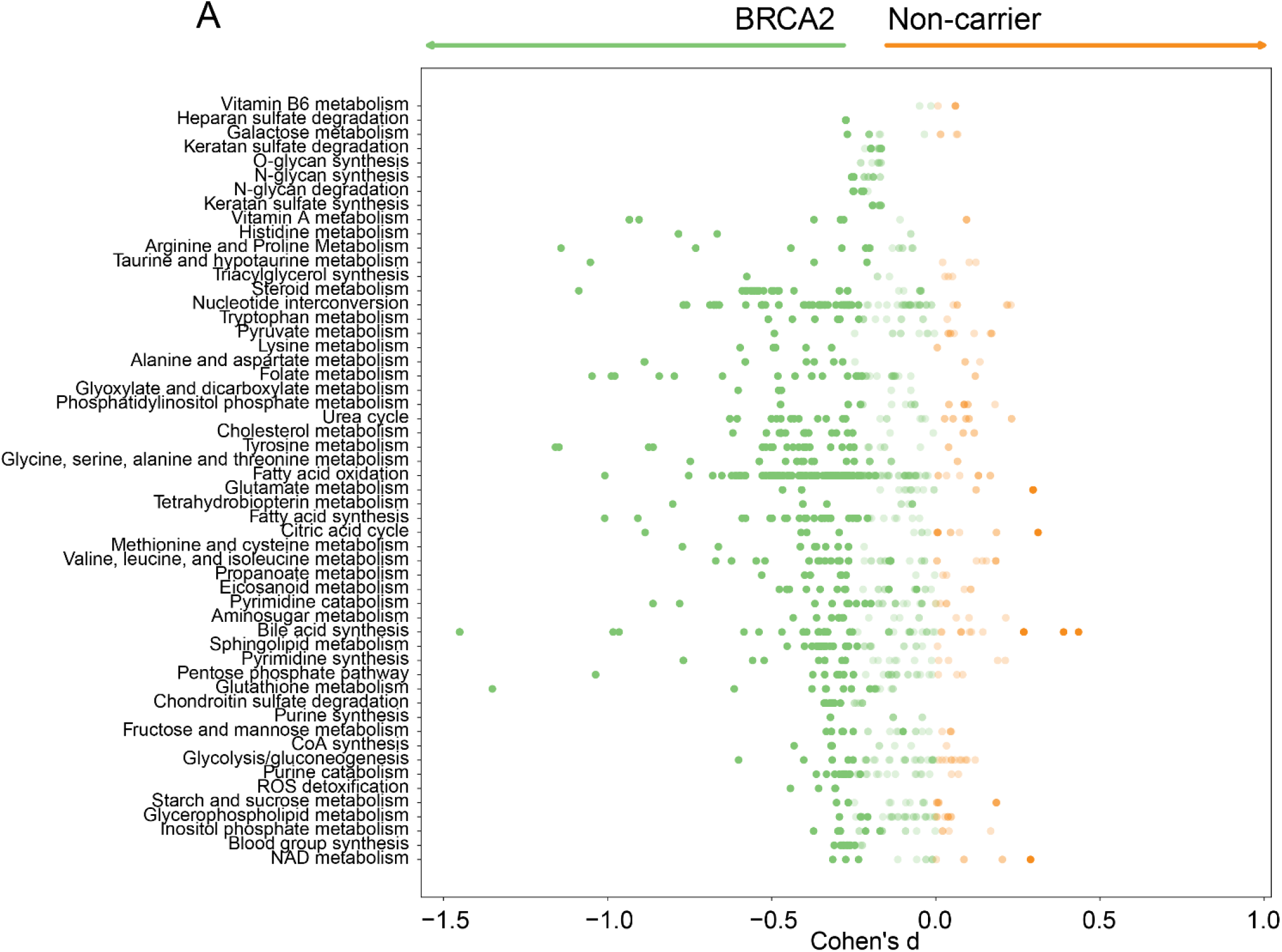
Metabolic subsystems in non-carriers vs. *BRCA2* FABP7 luminal progenitors (n = 294 cells per group).

**Figure S7:**
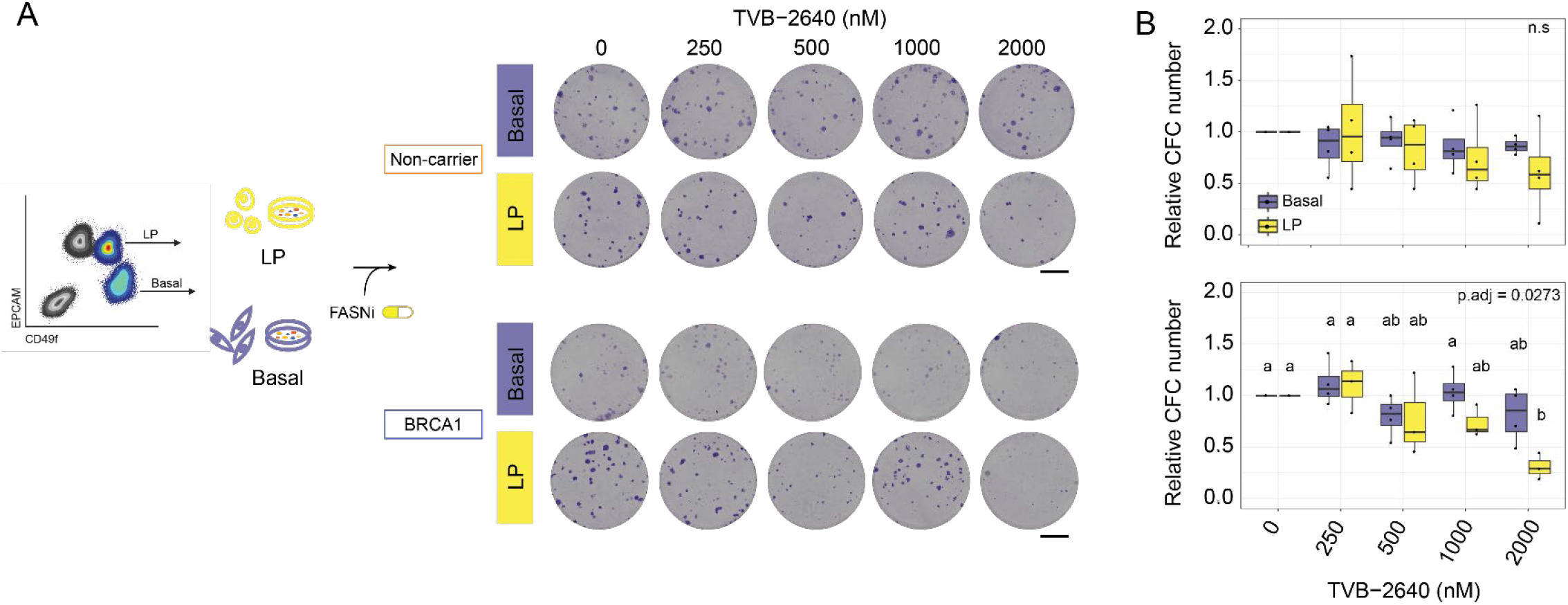
FASN inhibition in total FACS-purified luminal progenitor (LP) and basal mammary epithelial cells in non-carrier and *BRCA1* carrier. (A) Sorting schematic for isolation of total LP and total basal cells from *BRCA1* and non-carriers (left), and representative CFC images at day 9 post-treatment with FASNi (TVB-2640). **(B)** Colony number relative to DMSO control 9 days after TVB-2640 treatment (n = 4 per cell fraction per genotype).

